# Epigenetic analysis of Paget’s disease of bone identifies differentially methylated loci that predict disease status

**DOI:** 10.1101/2021.01.04.425216

**Authors:** Ilhame Diboun, Sachin Wani, Stuart H Ralston, Omar M E Albagha

## Abstract

Paget’s Disease of Bone (PDB) is characterized by focal increases in disorganized bone remodeling. This study aims to characterize PDB associated changes in DNA methylation profiles in patients’ blood. Meta-analysis of data from the discovery and replication set, comprising of 116 PDB cases and 130 controls, revealed significant differences in DNA methylation at 14 CpG sites, 4 CpG islands, and 6 gene-body regions. These loci, including two characterized as functional through eQTM analysis, were associated with functions related to osteoclast differentiation, mechanical loading, immune function, and viral infection. A multivariate classifier based on discovery samples was found to discriminate PDB cases and controls from the replication with a sensitivity of 0.84, specificity of 0.81, and an area under curve of 92.8%. In conclusion, this study has shown for the first time that epigenetic factors contribute to the pathogenesis of PDB and may offer diagnostic markers for prediction of the disease.

## Introduction

Paget’s disease of bone (PDB) is characterized by increased but disorganized bone remodeling, which causes affected bones to enlarge, become weak and deform. The axial skeleton is predominantly involved, and commonly affected sites include the skull, spine, pelvis, femora and tibia. Paget’s disease is clinically silent until it has reached an advanced stage at which point irreversible damage to the skeleton has occurred (1). Bisphosphonates are an effective treatment (2) and can often improve bone pain but have a limited impact on other clinical outcomes in patients with advanced disease (3, 4). On a cellular level, PDB is characterized by increased osteoclast activity and biopsies from affected bone lesions exhibit increase in the number and size of osteoclasts.

Genetic factors play an important role in classical PDB and in monogenic PDB-like syndromes (5, 6). Mutations in *SQSTM1* are the most common cause of PDB but other susceptibility genes and loci have been identified through genome wide association studies (7–9). These include genes that play an important role in osteoclast differentiation such as *CSF1*, *TNFRSF11A* and *DCSTAMP*. Additionally, an expression quantitative trait locus (eQTL) in *OPTN* is associated with increased susceptibility to PDB (10). Functional analysis using mouse models showed that OPTN is a negative regulator of osteoclast differentiation and mice with loss of *OPTN* function develop PDB-like bone lesions with increasing age (10, 11).

Environmental factors also play a role, as evidenced by the fact that the disease is focal in nature and its incidence and severity has diminished in recent years (12). Several environmental triggers have been suggested including persistent viral infection, repetitive mechanical loading of the skeleton, low dietary calcium intake, environment pollutants and vitamin D deficiency (6).

The possible role of persistent viral infection with measles and distemper has been studied experimentally. for example, expression of the measles virus nucleocapsid protein in osteoclasts was found to trigger PDB-like phenotype in mice (13, 14). However, clinical studies which have sought to detect evidence of viral proteins and nucleic acids in humans with PDB have yielded conflicting results (2).

Accumulating evidence suggests that environmental and lifestyle factors can influence gene expression and clinical phenotype in various diseases through epigenetic mechanisms such as changes in DNA methylation. To gain insights into the role of epigenetic DNA methylation in PDB, we have conducted genome-wide profiling of DNA methylation in a cohort of 253 PDB patients and 280 controls and evaluated the predictive role of epigenetic markers in differentiating patients with PDB from controls.

## Results

### Characteristics of study cohort

Table 1 shows descriptive statistics for the study cohort. PDB cases in the discovery set were slightly older and included more males compared to controls but no difference in age or gender distribution was found in the replication set. The number of patients with *SQSTM1* mutations was similar in the discovery and replication set and accounts for approximately 14% of PDB cases.

**Table 1.** Descriptive statistics of the study cohort

|  | Discovery |  | Replication |  |
| --- | --- | --- | --- | --- |
|  | PDB Case | Control | PDB Case | Control |
| Number | 116 | 130 | 116 | 130 |
| Age (years), mean ± SD | 72.1 ± 7.5* | 70.0 ± 7.4* | 72.5 ± 8.7 | 72.3 ± 8.2 |
| Males, n (%) | 65 (56.0)* | 48 (36.9)* | 59 (50.9) | 53 (40.8) |
| Females, n (%) | 51 (44.0)* | 82 (63.1)* | 57 (49.1) | 77 (59.2) |
| *SQSTM1* Mutation n (%) | 16 (13.8) | 0 (0) | 17 (14.6) | 0 (0) |

\*P<0.05 comparing Paget's disease (PDB) cases to controls.

### Differentially methylated Sites (DMS)

Figure 1 shows the study design and summary of differential methylation results. After adjusting for all confounders, differential methylation analysis of the discovery set revealed 419 DMS with FDR < 0.05, 57 of which reached statistical significance (FDR < 0.05) in the replication set (Table S1). Meta-analysis of the DMS from discovery and replication revealed 14 Bonferroni significant DMS (P< 1.17 × 10^−7^; Table 2). The direction of effect for all replicated DMS was identical in the discovery and replication set and shows hypermethylation in PDB cases compared to controls. A Manhattan plot of the results is shown in Figure 2-A.

**Table 2.** Differentially methylated CpG sites (DMS) in Paget’s disease of bone

| CpG Site |  |  | Discovery |  | Replication |  | Meta-analysis |  | Annotations |
| --- | --- | --- | --- | --- | --- | --- | --- | --- | --- |
| Probe ID | Chr | Position | $\Delta$ Beta* | P Value | $\Delta$ Beta* | P Value | $\Delta$ Beta* | P Value | Nearest gene |
| cg10290814 | 17 | 7284330 | -0.018 | 1.2x10<sup>-6</sup> | -0.015 | 1.4x10<sup>-4</sup> | -0.017 | 2.3x10<sup>-10</sup> | *TNK1* |
| cg19361865 | 1 | 220922163 | -0.014 | 5.4x10<sup>-6</sup> | -0.012 | 9.7x10<sup>-5</sup> | -0.013 | 7.6x10<sup>-10</sup> | *MOSC2* |
| cg09152582 | 1 | 88928362 | -0.021 | 2.1x10<sup>-5</sup> | -0.018 | 3.5x10<sup>-5</sup> | -0.019 | 1.1x10<sup>-9</sup> | *PKN2-AS1* |
| cg09260089 | 10 | 134599860 | -0.024 | 4.6x10<sup>-5</sup> | -0.024 | 1.2x10<sup>-4</sup> | -0.024 | 9.5x10<sup>-9</sup> | *NKX6-2* |
| cg24879273 | 10 | 102989645 | -0.026 | 4.9x10<sup>-5</sup> | -0.016 | 1.7x10<sup>-4</sup> | -0.021 | 1.4x10<sup>-8</sup> | *LBX1* |
| cg03839709 | 13 | 96743492 | -0.014 | 2.7x10<sup>-4</sup> | -0.014 | 3.4x10<sup>-5</sup> | -0.014 | 1.8x10<sup>-8</sup> | *HS6ST3* |
| cg16419235 | 8 | 57360613 | -0.036 | 1.9x10<sup>-4</sup> | -0.029 | 8.3x10<sup>-5</sup> | -0.032 | 3.1x10<sup>-8</sup> | *PENK* |
| cg04317962 | 16 | 79623625 | -0.017 | 1.4x10<sup>-6</sup> | -0.019 | 2.9x10<sup>-3</sup> | -0.018 | 3.1x10<sup>-8</sup> | *MAF* |
| cg01429039 | 4 | 52918065 | -0.023 | 1.8x10<sup>-4</sup> | -0.020 | 1.1x10<sup>-4</sup> | -0.021 | 3.5x10<sup>-8</sup> | *SPATA18* |
| cg03885399 | 1 | 47691550 | -0.020 | 4.4x10<sup>-6</sup> | -0.014 | 3.6x10<sup>-3</sup> | -0.017 | 4.7x10<sup>-8</sup> | *TAL1* |
| cg04738965 | 3 | 147127662 | -0.037 | 4.0x10<sup>-5</sup> | -0.028 | 7.1x10<sup>-4</sup> | -0.033 | 6.2x10<sup>-8</sup> | *ZIC1* |
| cg10954182 | 12 | 104532377 | -0.016 | 1.9x10<sup>-4</sup> | -0.009 | 2.1x10<sup>-4</sup> | -0.013 | 7.8x10<sup>-8</sup> | *NFYB* |
| cg10964367 | 8 | 1771973 | -0.025 | 1.3x10<sup>-4</sup> | -0.019 | 3.8x10<sup>-4</sup> | -0.022 | 9.4x10<sup>-8</sup> | *ARHGEF10* |
| cg12739454 | 1 | 164290833 | -0.018 | 2.4x10<sup>-4</sup> | -0.012 | 2.4x10<sup>-4</sup> | -0.015 | 1.1x10<sup>-7</sup> | - |

\* $\Delta$ Beta represents the difference in DNA methylation in cases as compared to controls (Beta Control-Beta PDB). Position in base pairs in reference to human genome build 37 (GRCh37). Chr, chromosome; CpG, cytosine-phosphate-guanine. All P-values are genome-wide significant based on Bonferroni corrected pvalue < 0.05.

**Figure 1.**
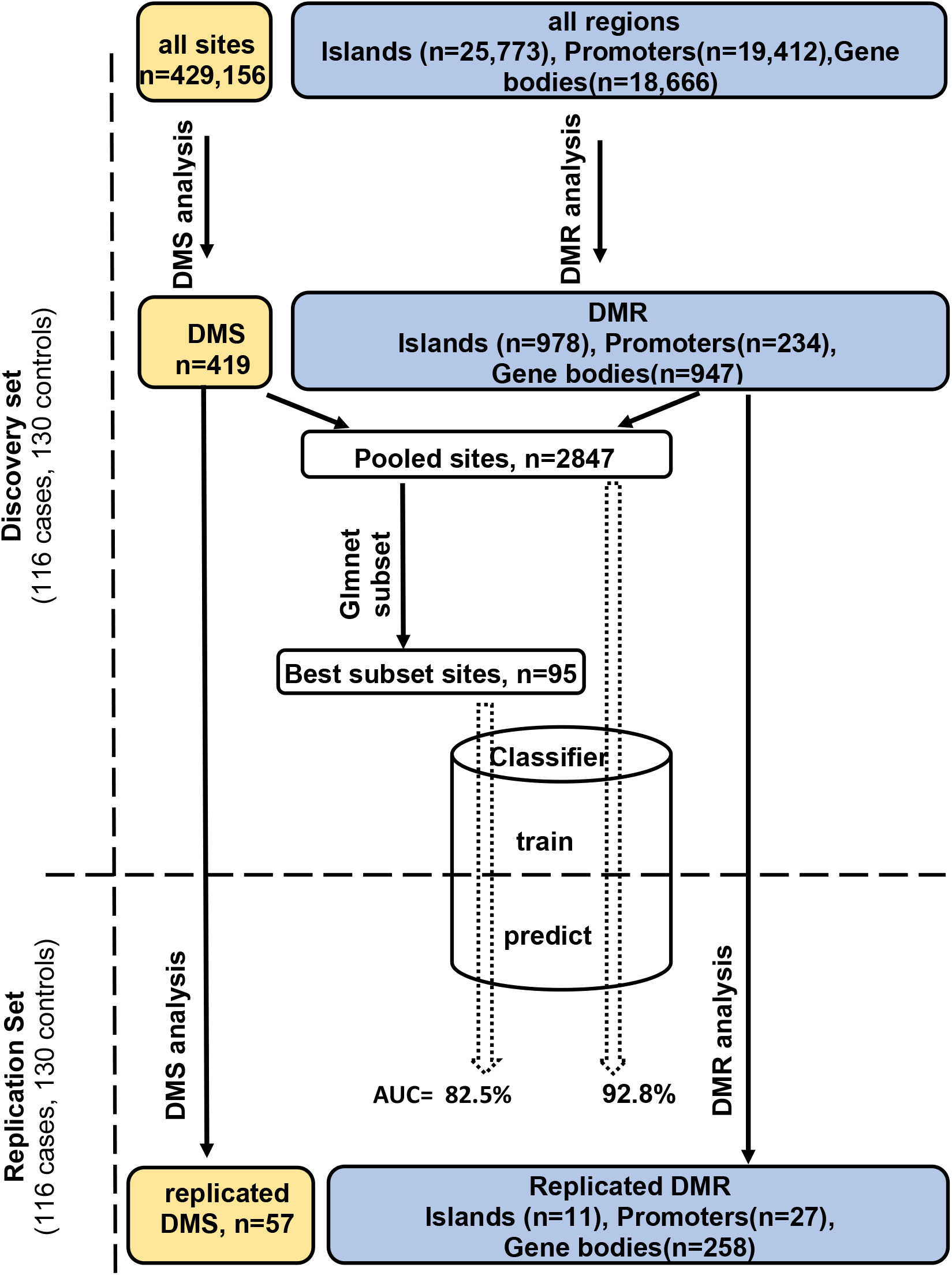
Study design and analysis workflow. Differentially methylated sites (*DMS*) and differentially methylated regions (*DMR*) were analyzed using, the General/Generalized linear model respectively, in the discovery set. Those reaching FDR < 0.05 were tested in the replication set to identify DMS/DMR that replicate at the same significance level. The DMS and the important sites within DMR were pooled together giving rise to the *Pooled sites* (refer to methods), of these a best PDB discriminatory subset was obtained using the Lasso and Elastic-Net regression. A multivariate classifier based on the discovery measurement of the Pooled/Best subset sites yielded an AUC value of 92.8% and 82.5% respectively when tested in the replication.

**Figure 2.**
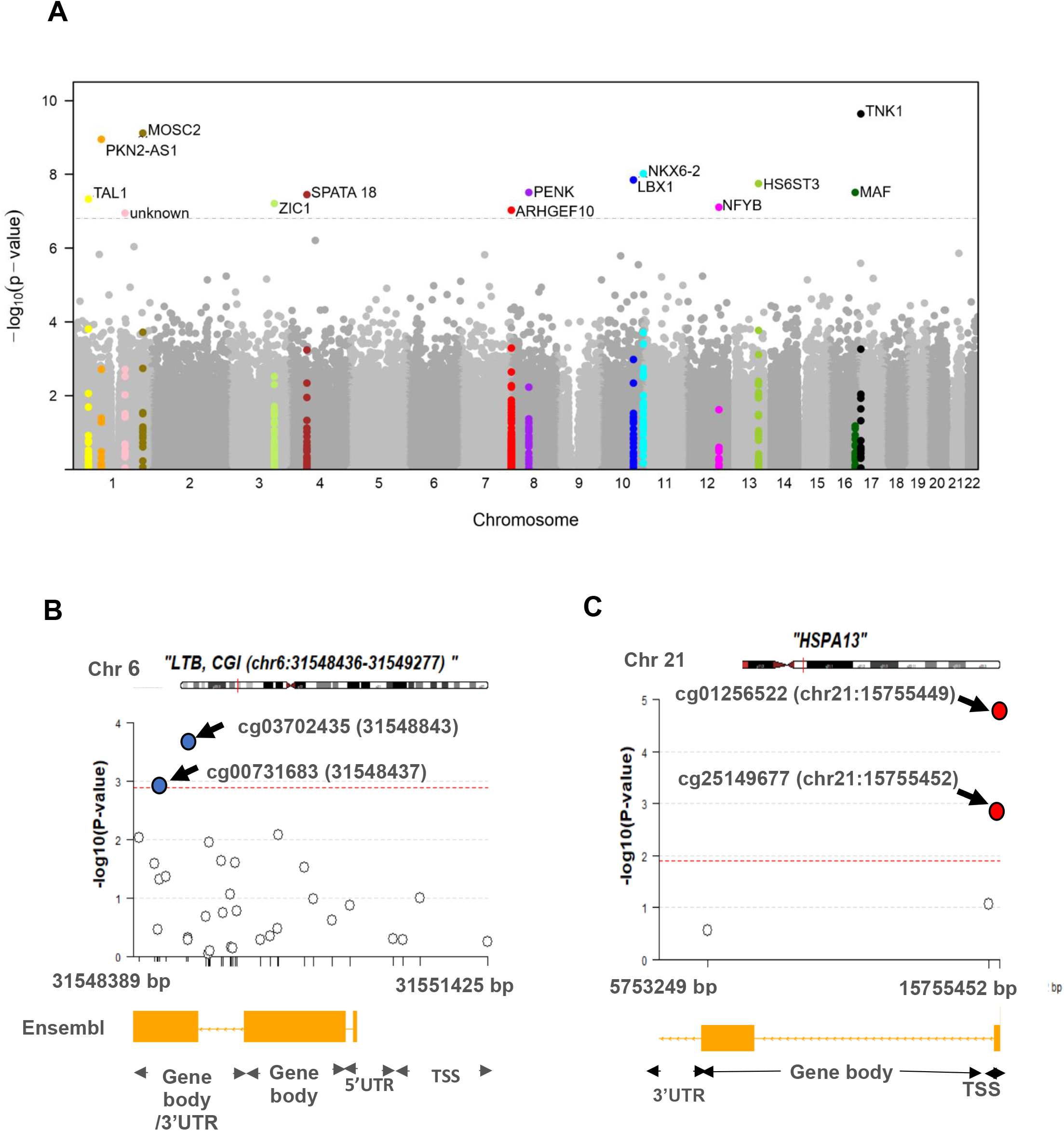
Differential methylation analysis comparing controls to PDB patients (n=246). A) Site analysis, a Manhattan plot showing the chromosomal positions (x-axis) versus the -log10 (P) of significant DMS and adjacent sites. For the Bonferroni significant sites however, the meta-analysis P-values are shown instead and highlighted in color. The horizontal dashed line indicates the Bonferroni corrected significance threshold (P< 1.17 × 10^−7^). B&C) Region analysis, showing the multitude of significantly hypermethylated (red) and hypomethylated (blue) sites from LTB (Bonferroni replicated from island analysis) and HSPA13 (Bonferroni replicated from gene body analysis). The dashed lines represent the fdr < 0.05 threshold for each region which depends on the number of sites within the region (refer to methods).

### Differentially methylated regions (DMR)

Besides analyzing individual sites, our region-based analysis was intended to uncover densely hyper/hypo-methylated regions across the genome in PDB as well as identifying instances where the effect from individual sites is moderate, yet accumulatively significant. We tested natural concentrations of sites within CpG islands but also gene bodies and promoter regions, justified by the fact that promoter methylation often suppresses transcription whilst that from the gene body often stimulates gene expression (Figure 1).

Evaluation of the 25,773 CpG islands on the array, revealed 978 DMR that were significantly differentially methylated (FDR < 0.05) in the discovery set, 111 of which replicated at the same significance level in the replication set (Table S2). Stringent Bonferroni multiple testing correction revealed 4 islands that remained significant following discovery and replication, and these were located near *LTB*, *SKIV2L*, *EBF3* and *CCND1* (Table 3).

**Table 3.**
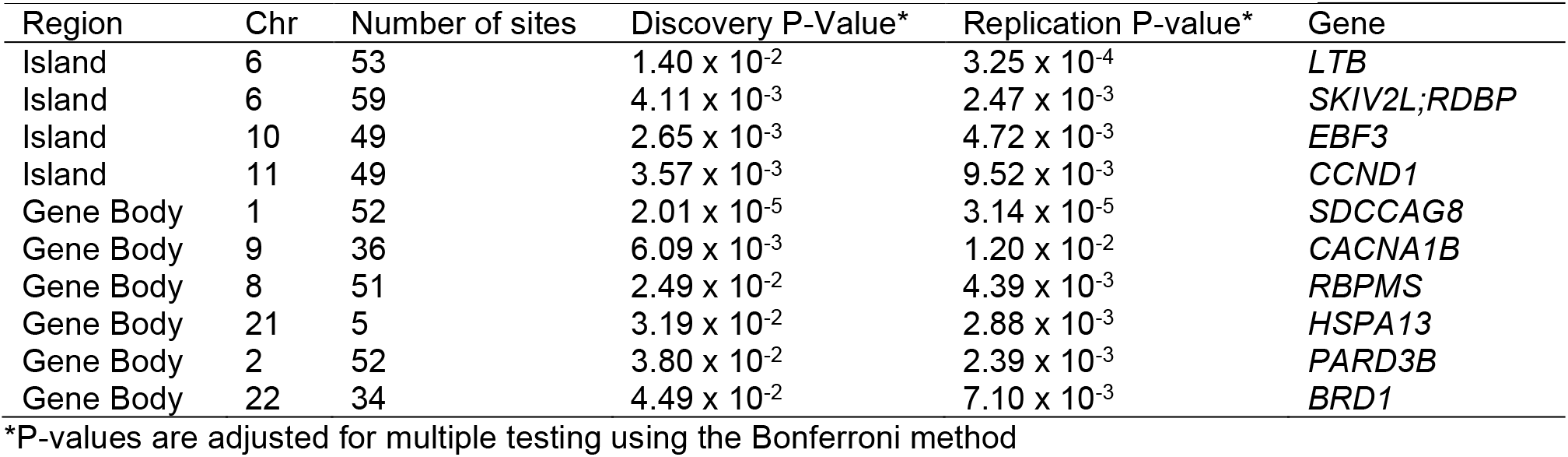
Differentially methylated regions (DMR) in Paget’s disease of bone

Gene body analysis revealed 258 (FDR < 0.05) replicated DMR out of a total of 947 differentially methylated genes initially identified in the discovery set (Table S3). Six gene body DMR reached significance after Bonferroni correction in both the discovery and replication set (Table 3). In the context of promoter regions, evidence for FDR significant association with the disease was equally observed in the discovery and replication set for 27 promoters DMR (Table S4), but none reached significance after Bonferroni correction. Figure 2-B&C show a regional plot for DMR LTB and HSPA13 from island and gene body analysis respectively, highlighting the co-occurrence of multiple differentially methylated sites along each region.

### Mapping common regulatory patterns of DNA methylation into functional networks

To gain further insight into the pathology of PDB, we explored common methylation patterns amongst functional keywords identified as significantly over-represented amongst the *Pooled sites.* Figure 3 shows a graphical representation of these functional networks. In addition to bone-related cells, there is a strong presence of immune cells linked to key biological processes including proliferation, differentiation, autophagy and cell death. Furthermore, virus, cytokines, and interferon-gamma were among the over-represented keywords. The process of ubiquitination lies at the center of the graph with the largest number of links in the network.

**Figure 3:**
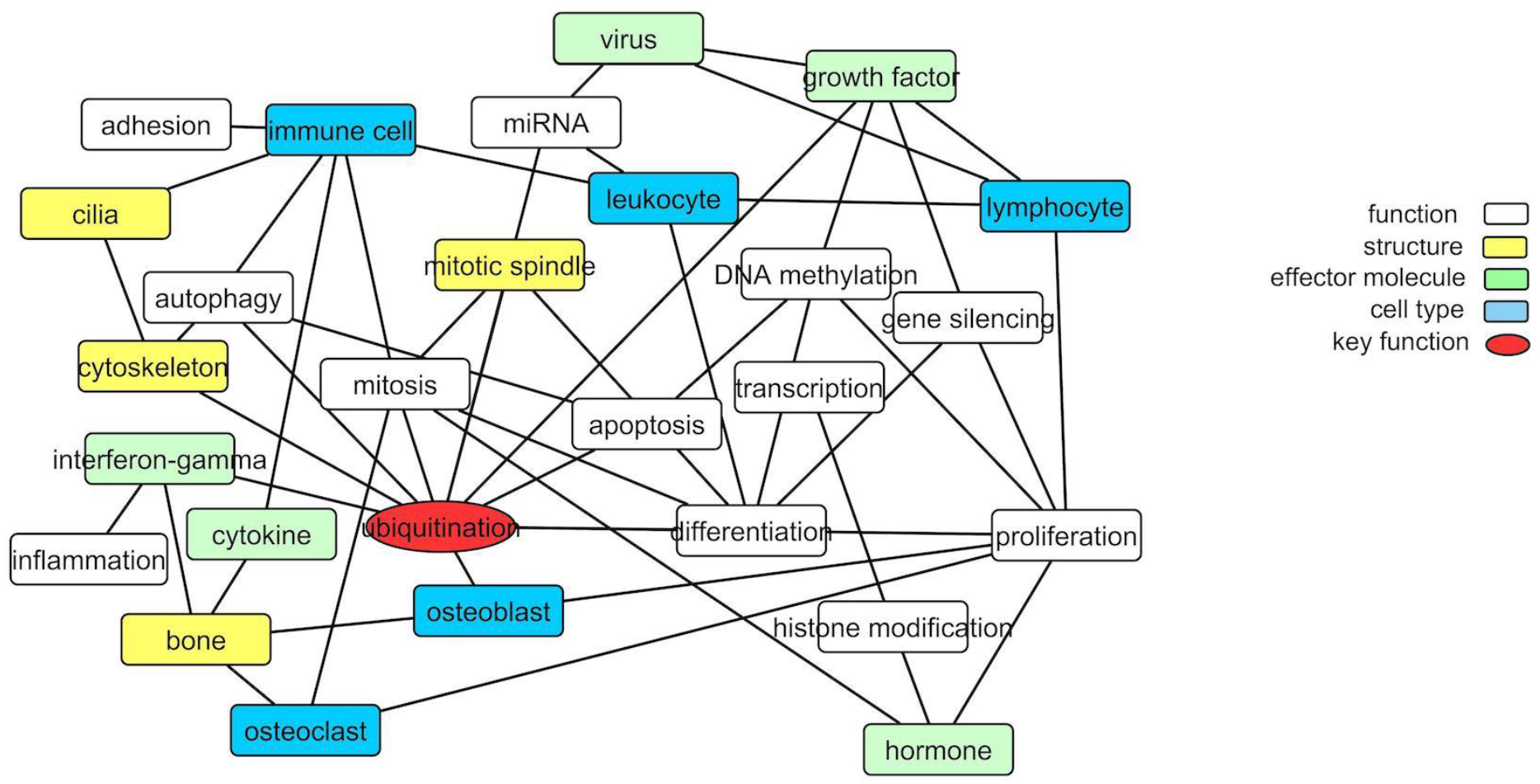
Translating the methylation data into functional network. Nodes are functional, cellular, molecular and sub-cellular keywords from GO annotations enriched amongst the *Pooled sites*. An edge between two nodes indicates that differentially methylated genes associated with the keyword in node 1 are significantly partially correlated with their counterparts from node 2 more often than can be accounted for by chance.

### Diagnostic capacity of differentially methylated markers

In order to determine whether differentially methylated markers might be of diagnostic value, we performed OPLS-DA in the discovery and replication cohorts. The results are summarised in Figure 4. The OPLS-DA procedure was first performed using the combined set of significant DMS and DMR identified from the discovery set (*Pooled sites*; n=2847, refer to methods for further details) and when the classifier was tested on the replication set, it yielded an AUC of 92.8%. To identify sites with the highest predictive ability, we applied the Net Regularization Extension of the Generalized Linear Model approach on the *Pooled* sites which highlighted 95 sites (*Best subset* sites; Table S5), out of the 2847 initial *Pooled* sites, as best discriminatory of PDB cases and controls (Figure 1). The OPLS-DA procedure performed on this *Best subset* resulted in an AUC of 82.5%. A rather superior performance in comparison to similarly trained classifiers based on the DMS (AUC=67%), islands DMR (AUC = 76%), or promoter DMR (AUC = 79%) analyses. On the other hand, the AUC from a classifier restricted to the DMR gene bodies was 92% which is similar to that obtained from the whole *Pooled sites* (AUC 92.8, Figure 3).

**Figure 4.**
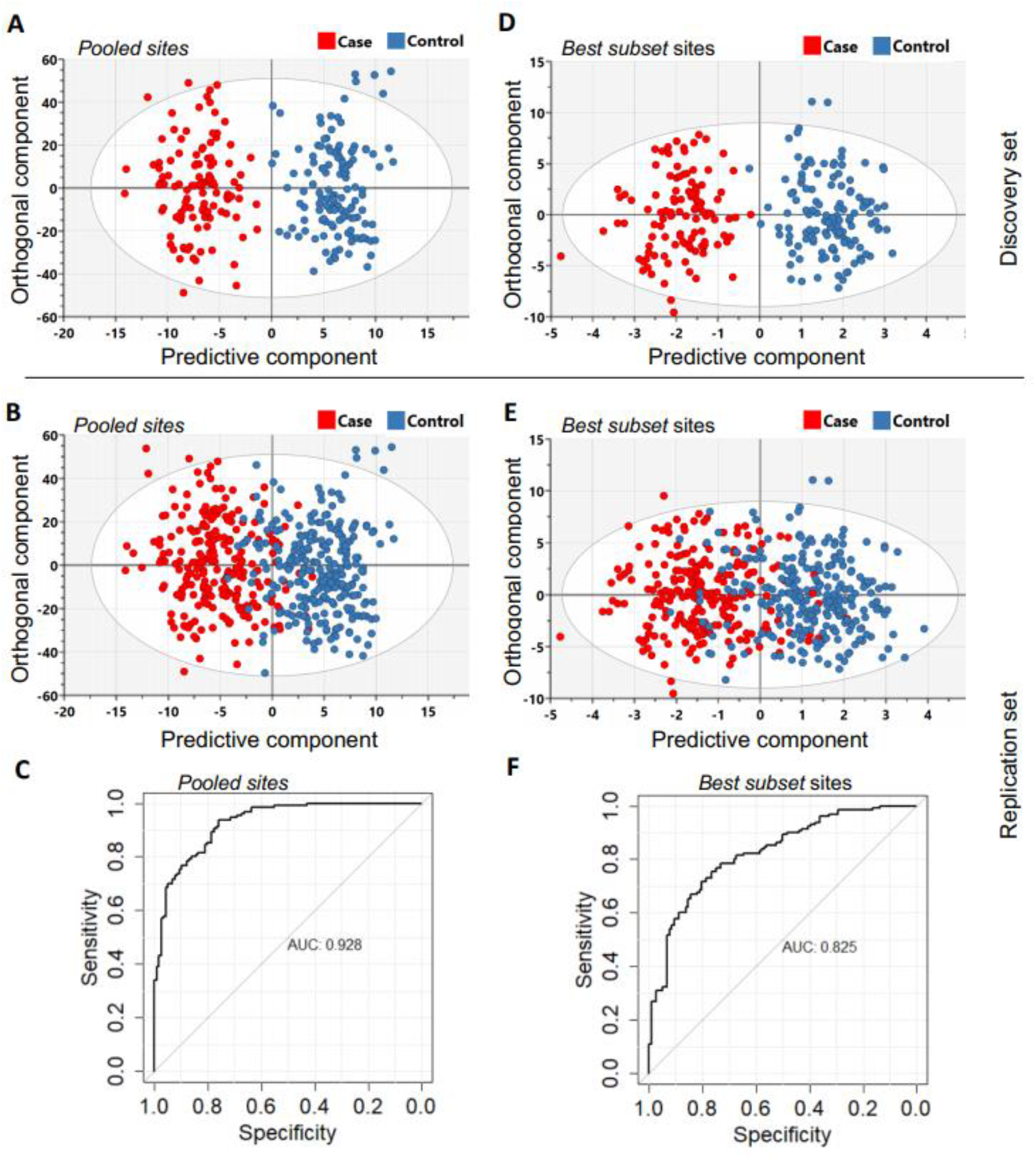
The orthogonal partial least squares discriminant analysis (OPLS-DA) was performed using the *Pooled sites* identified from the discovery set (n=246). (**A**) Classifier trained on all 2847 pooled sites with FDR < 0.05 (*Pooled sites*) from the discovery set. (**B**) Testing the classifier on the replication set. (**C**) ROC curve analysis yielded an overall sensitivity of 0.84, specificity of 0.81 and AUC=0.928. (**D**) Classifier trained on the *Best subset* sites from Glmnet analysis (n=95) using the discovery set. (**E**) Testing the classifier on the replication set. (**F**) ROC curve analysis showed an overall sensitivity of 0.77, specificity of 0.74 and AUC=0.825. The Scatter plots in A,B,C&D show the predictive component that discriminates PDB cases from controls (x-axis) versus the orthogonal component representing a multivariate confounding effect that is independent of PDB (y-axis).

Functional enrichment analysis of the 95 *Best subset* was consistent between IPA and GO with many genes annotated to the following broad functional terms: *immune function*; *bone lesions and bone homeostasis,* and *viral processes*. Several identified genes fell into more than one category. Overlaying the IPA knowledge-based repository of molecular interactions identified a handful of functional links between the genes located in the *Best subset* sites, highlighting important functional subnetworks (Figure 5-A). Additionally, we found that the effect size (absolute difference in DNA methylation between controls and PDB cases) was significantly higher for sites from the *Best subset* (mean ± SD; 0.011 ± 0.019) compared to the rest of those in the *Pooled sites* (0.007 ± 0.01; P-value = 1.9 × 10^−3^). The magnitude of effect from each site in the *Best subset,* as calculated by the Elastic-Net Regularization Extension of the Generalized Linear Model, is color-coded in Figure 5-B.

**Figure 5:**
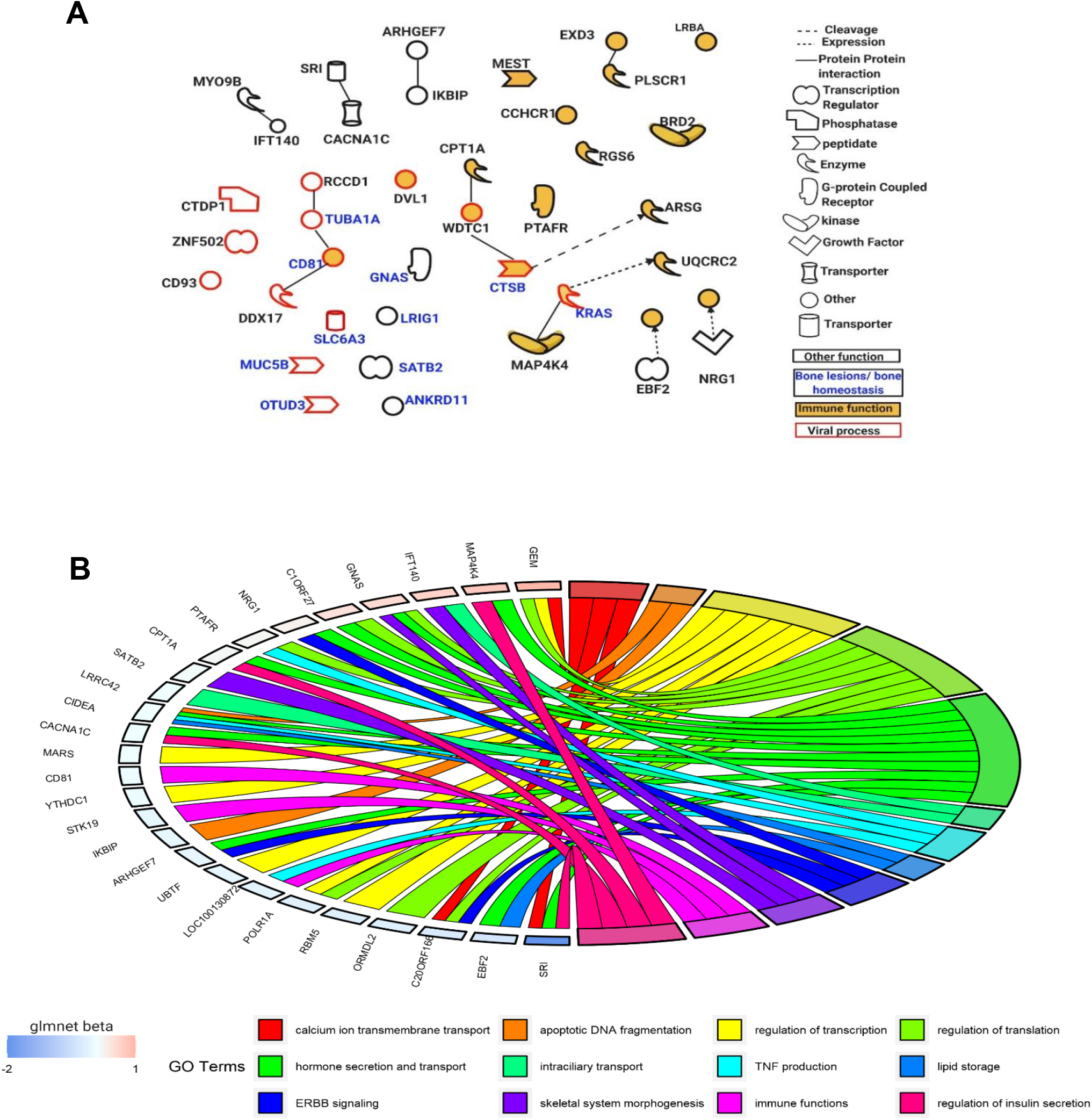
Functions of genes mapped near the *Best subset* of differentially methylated sites identified through. the Elastic-Net Regularization Extension of the Generalized Linear Model. A) An IPA based network showning a subset of these genes with functional interactions (edges) or mapping to one of three functional classes: immune, viral and bone homeostasis. B) An overview of GO biological processes significantly enriched amongst the Best subset together with their beta values from the Glmnet R package implementing the extended Generalized Linear Model in question.

### eQTM analysis

eQTM analysis showed that the the Bonferroni significant DMS cg10964367 was associated with the expression level of ARHGEF10 (P = 3.9 × 10^−9^). Additionally, cg26724726 from gene body analysis was associated with the expression of LTB (P = 1.10 × 10^−5^) and 8 of the *Best subset* sites were associated with the expression of nearby genes (Table S6).

## Discussion

The present study is the first to investigate DNA methylation profiles in Paget’s disease of bone. DNA methylation profiles from PDB patients were compared to controls and Meta-analysis of discovery and replication revealed 14 genome-wide significant DMS. Many were located within or near genes with functional relevance to the pathogenesis of PDB including bone-related functions, such as osteoclast differentiation, or functions related to environmental triggers associated with PDB such as viral infection and mechanical loading. TNK1 is a tyrosine kinase that has a pivotal role in innate immune responses by regulating the Interferon-stimulated genes downstream of the JAK-STAT pathway (15). It has previously been associated with frontotemporal dementia (16) which can co-exist with Paget’s disease (17). MOSC2 is a member of the membrane-bound E3 ubiquitin ligase family that regulates endosome trafficking (18) Less is known about the specific functions of transcription factors NKX6-2 and LBX1 in bone metabolism, but mutations in the latter are associated with Scoliosis HS6ST3 plays a key role in the synthesis of heparan sulfate that potentiates key growth factors including the bone morphogenic protein BMP and Wnt (19). PENK encodes for proenkephalin, the precursor of a range of effector molecules including pain-associated pentapeptide opioids as well as modulators of osteoblast differentiation (20). Interestingly, PENK knockout mice have abnormal bone structure and mineralization (21). MAF was found to promote osteoblast differentiation and heterozygous deletion of MAF in mice results in age-related bone loss associated with accelerated formation of fatty marrow (22). SPATA18 is expressed in a variety of cancers including osteosarcoma and its transcription is induced by p53 (23). TAL1 has been found to regulate osteoclast differentiation through suppression of their fusion mediator DCSTAMP (24). The Zinc finger protein ZIC1 has a role in shear flow mechanotransduction in osteocytes (25). Expression of ZIC1 in human was found to be increased in loaded compared to unloaded bone and the increased expression in loaded bone is associated with reduced methylation in several CpGs in ZIC1 (26). NFYB confers chromatin access to other transcriptional regulators and is known to be involved in transition through cell cycle (27). Finally, the centrosomal ARHGEF10 has a role in the formation of mitotic spindle during mitosis (28).

Our analysis was extended to identify regions with frequent methylation changes in PDB amongst adjacent sites. Genomic regions have traditionally been evaluated in epigenetics studies based on linear combinations of methylation data from residing sites or through meta-analysis of effects/p-values from an initial site-level differential methylation analysis. The novel approach presented in this study is advantegous in to ways: First, our method allows for sites to be hyper or hypo methylated along the same region unlike the linear combination approach where opposing effects could neutralize one another. Second, it draws strength from the collective effects of neighbooring sites whilst avoiding the limitations of the site-level analysis approach.

Four Bonferroni significant DMR were identified in islands which were located near the following genes: LTB, a cytokine shown to simulate osteoclast activity (29); SKIV2L, with an RNA helicase activity, thought to be involved in blocking translation of viral mRNA and has been implicated in regulating host responses to viral infections (30); EBF3 which is involved in bone development and B cell differentiation (31) and CCND1, a Wnt target that was reported to be upregulated in response to mechanical loading of bone (32).

Additionally, six Bonferroni significant DMR in gene bodies were identified. These were located within genes with functions related to mitosis and ciliogenesis (SDCCAG8) (33); TGFB1-mediated signaling (RBPMS) (34); calcium signaling (CACNA1B) (35); protein ubiquitination (HSPA13) (36); cytoskeletal organization (PARD3B) (37) and histone acetylation (BRD1) (38).

The *Pooled* sites identified from the discovery set were able to discriminate cases and controls with a considerable accuracy when tested on the replication set. The Best subset analysis allowed the identification of a smaller subset of sites trading off the classification accuracy with the number of explanatory sites. The AUC of 82.5%, based on the 95 discriminatory sites from the best subset analysis, is promising and future experiments are warranted to study its clinical applicability.

In terms of disease pathology, the DNA methylation data reflected many environmental triggers thought to be involved in PDB. Some of the genes amongst the DMS and the 95 *Best* subset were associated with immune antiviral responses (Figure 5 & Table S5). This is of interest since a previous study in the PRISM cohort showed that levels of antibodies to Mumps virus were significantly higher in PDB cases compared to controls (39). Although we and others have failed to detect evidence of ongoing virus infection in PDB, the above data is consistent with the hypothesis that host immune responses to infection may be altered in PDB.

Differential methylation of ZIC1 and CCND1 indicate possible differences between cases and controls in these genes which are involved in mechano-transduction, a process which has been implicated in localisation of bone lesions in PDB (39, 40). Our study also highlighted genes that regulate the cell cycle, vesicular transport and cytoskeletal reorganization as being potentially involved in PDB. Other genes were identified that play a role in immune cell function and these were strongly represented in the best subset of differentially methylated sites. This lends support to the hypothesis that PDB may be a disorder with an osteoimmunological basis (41) and should prompt further work to investigate host-environment interactions including studies of the microbiome in this complex but fascinating disease (42).

Apart from providing new insights into the potential links between genes and environment in regulating susceptibility to PDB, this study has revealed the potential role of methylation signals as a biomarker for disease susceptibility. Potent bisphosphonates such as zoledronic acid can return the abnormalities of bone remodeling to normal in a large proportion of patients with PDB (4, 43). Unfortunately PDB often remains clinically silent until it has reached an advanced stage by which point irreversible skeletal damage may already have occurred (5). This study raises the possibility that epigenetic markers, possibly when combined with genetic profiling would be worth exploring as means of assessing the risk of developing PDB in people with a family history of the disorder so that early intervention can be considered where clinically appropriate.

One limitation of the study is the fact that the identified methylation changes were not shown to occur in the osteoclasts which are the cells of main interest in Paget’s pathogenesis. This is primarily justified by the difficulty to collect bone tissue from PDB patients in a similarly sized cohort. Moreover, showing an epigenetic signature to PDB in the blood adds to the increasing evidence in the literature pointing to the possibility of pathogenic immune processes lying at the heart of PDB. More importantly, a predictive epigenetic signature in a readily accessible tissue such as the blood has clinical implication, also considering the silent nature of PDB and the possibility of avoiding much of the adverse symptoms of the disease with early diagnosis. Finally, one needs to consider that blood also contains progenitors of bone cells and that white blood cells share a similar ancestry with osteoclasts.

## Materials and methods

### Study Subjects

The DNA samples were derived from UK-based PDB patients and controls who took part in the PRISM trial (Paget’s Disease: Randomized Trial of Intensive versus Symptomatic Management (ISRCTN12989577) (44). The PRSIM trial is a multi-center study in which participants were recruited from 27 different clinical centers across the United Kingdom. The epigenetic analysis was conducted in 253 cases with clinical and radiological evidence of PDB and 280 controls who were spouses of PDB cases (n=135) or subjects who had been referred for investigation of osteoporosis but had normal bone density upon examination by dual energy X-ray absorptiometry (n=131). The cohort was randomly divided into a discovery and replication set comprising of comparable numbers of cases and controls (Figure 1). According to the study by Tsai and Bell, a 10% difference in the mean of CpG methylation level between cases and controls at genome-wide significance level of 10^−6^ requires 112 individuals in each group to achieve 80% EWAS power (45). On this basis, our discovery set comprising of 116 cases and 130 controls is adequately powered and the results are further validated in an equally sized replication set.

### DNA methylation profiling

Genomic DNA was extracted from peripheral blood using standard protocols. Bisulfite conversion was performed on 500μg of DNA using Zymo EZ-96 DNA methylation Kit (Zymo Research, USA). DNA methylation profiling was performed using the Illumina Infinium HumanMethylation 450K array (Illumina, USA) by following the manufacturer’s protocol. The R package *RnBeads* version 1.10.8 was used for quality control (8). Samples with low methylated or unmethylated median intensity (<11.0) were excluded (n=35) along with samples with sex mismatch between reported and predicted sex (n=0). Probes with the following criteria were excluded: detection P value >0.05, cross reactive probes, containing a SNP within 3 bp of nucleotide extension site, or those located on sex chromosomes. Additionally, 723 sites were further excluded from the dataset for previously established association with smoking (46). A total of 56,356 probes were excluded from the initial 485,512 leaving 429,156 CpGs for analysis (Figure 1). The final dataset used for analysis comprised of 232 PDB cases and 260 controls. The *Enmix* method (47) was used for background correction whilst *SWAN* was used to achieve between and within array normalization. For all downstream analysis, the M-values, derived using the formulae log2((methylated signal+1)/(unmethylated signal+1)), were used.

### Statistics

An overview of the analysis performed in this study is shown in Figure 1, in what follows we provide details of each analysis step:

#### Differential methylation analysis of sites

In order to account for the heterogenous cellular composition of the measured samples, the counts of the following cell types CD14 monocytes, CD19 B-cells, CD4 T-cells, CD56 NK cells, CD8 T-cells, eosinophils, granulocytes and neutrophils were estimated using the *Houseman* reference method (48), part of the *RnBeads* pipeline. The reference methylome was obtained from previously published methylation data measured from sorted blood cells comprising 47 samples (49). These reference samples were normalized together with our data to make sure that extrapolation of cell type information was unaffected by differences between the two datasets. we performed Surrogate variable analysis (SVA) which captures additional unknown sources of variation based on joint methylation patterns amongst the different sites that do not correlate with the disease. The top 10 significant SVA components were extracted from the data using the SVA functionality in *RnBeads.*

In all statistical models described below, the term *confounders* refer to the following covariates: age, sex, array, bisulfite conversion batch, array scan batch, blood cell composition from the *Houseman* method(48) and the top 10 surrogate variant analysis (SVA) components. The term *phenotype* denotes the control/PDB state of each sample. The term *region* is used to describe clusters of sites along the genome including CpG islands, gene bodies and promoters. CpG islands were delineated in the illumina array manifest file as well as RnBeads annotation libraries. Gene bodies and promoters were manually assigned. More specifically, sites mapping to the transcription start site (TSS) according to the manifest were attributed to a promoter region whilst those falling at the 5’ untranslated region or gene body were assigned to a gene body region. A general linear model based on the limma moderated standard error (50) was used to assess differentially methylated sites (DMS) between cases and controls using the model: *CpG site ~ phenotype + confounders*. The model was first run on all sites in the discovery set and all DMS with a significant FDR (< 0.05) in the discovery set were assessed in the replication set. Meta-analysis looking at the combined effect from both discovery and replication was performed on this subset using the R package *Metafor* (51). The Bonferroni adjusted genome wide significance threshold of P=1.17 × 10^−7^ (0.05/429,156) was used.

#### Differential methylation analysis of regions

Differentially methylated regions (DMR) were analyzed using binomial regression, member of the family of the generalized linear models, in two steps:

First the parameters of the *null* model, excluding the sites, were estimated as follows:

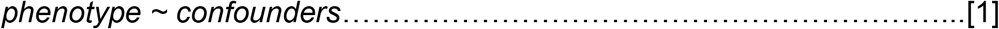

Next, all *n* sites within a given region (island/gene body/promoter) were incorporated into the model as follows:

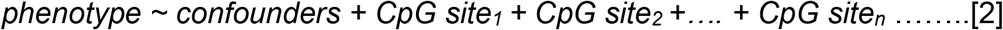

The difference in the deviance (equivalent to the residuals in the linear model) between the null model [1] and the full model [2] follows a χ^2^ distribution with n degrees of freedom. A P-value for the effect of the region given n sites was calculated accordingly. The analysis effectively tests for the significance of improvement in the model fit with the addition of the methylation data from the region of interest. The generalized linear model outlined above was run initially on the discovery set. The model was then repeated on the replication set on regions that were significant in the discovery set at FDR < 0.05. A similar approach was used to derive the Bonferroni significant regions. Visualization of the effect of individual sites from selected DMR was conducted using R package coMET (52).

#### Consolidating the DMS and DMR

In the Generalized Linear Model for region effect outlined in model formulae [2], the beta values from the individual sites are indicative of the sites’ level of association with the phenotype. This is effectively similar to the General Linear Model used for site-level analysis but with the important discrepancy that each site is being assessed while accounting for possible contributions of neighboring sites to the global effect of the region. We therefore extracted all the beta values form the full model in [2] from all the DMR. We then applied fdr based multiple testing correction on the pvalues corresponding to these beta values from fitting the model in [2] for each selected DMR separately. Sites with fdr < 0.05 were pooled with the DMS to create a unified list of significantly methylated sites or *Pooled sites* (Figure 1).

#### Discriminant analysis

Discriminant analysis was performed to assess the ability of the *Pooled sites* to tell apart cases from controls. We also used the Elastic-Net Regularization Extension of the Generalized Linear Model, provided by the R package *Glmnet (53)*, to identify the best subset of discriminatory sites (designated *Best subset*) of the list of *Pooled sites*. We trained an Orthogonal Projection to Latent Structures Discriminant Analysis (OPLS-DA) classifier (54), implemented in the software SIMCA ver. 15 (Umetrics, Sweden), on the discovery data from *Pooled* and *Best subset* sites separately. Each model was then tested on the replication set and its performance was further assessed based on the area under curve (AUC) value from receiver operating characteristic (ROC) curve analysis. The sensitivity and specificity measures of the test were estimated based on a classification threshold equal to the median of the predicted scores by the OPLS-DA classifier. The *Best subset* sites were analyzed further to reveal enrichment in biological functions. This was conducted using Ingenuity Pathway Analysis (IPA) (Qiagen, Germany) as well as the Gene Ontology (GO) R package *topGO* (55) based on the Fisher’s exact test statistics.

#### Partial correlation analysis of *Pooled sites*

Correlations in methylation patterns between CpG sites hold valuable information about how different biological functions are linked together in PDB. To this end, partial correlations between the *Pooled sites* were derived using the R package ggm (56). In parallel, the extensive GO functional annotations enriched amongst the genes associated with the *Pooled sites* were manually reduced to a manageable, yet representative, set of keywords: For instance, GO categories ‘regulation of proliferation’, ‘positive regulation of proliferation’ and ‘negative regulation of proliferation’ were all reduced to ‘proliferation’. The fisher’s exact test statistics was then used to assess whether the *Pooled sites* associated with a given keyword were correlated (based on the ggms) with their counterparts from another functional keyword more often than can be accounted for by chance alone. Fisher’s test p-values < 0.05 after FDR multiple testing correction were used to create pairs of functionally related keywords. The software Cytoscape (57) was used to visualize these associations.

### Expression quantitative trait methylation (eQTM) analysis

To assess the effect of DNA methylation at CpGs sites on the expression of nearby genes, we used data from the BIOS QTL browser (58).

## Acknowledgements

We wish to thank the patients and controls from the different centers who agreed to participate in this study. We would like to thank members of the PRISM trial research group across all participating centers for making DNA samples and data available for this study. We thank the Wellcome Trust Clinical Research Facility at Edinburgh University for performing the DNA methylation profiling.

## Competing interests

Prof S H Ralston has received research funding from Amgen, Eli Lilly, Novartis, and Pfizer unrelated to the submitted work. The other authors have no conflicts of interest to declare.

## Notes

**Funding** This work was funded by a consolidator grant from the European Research Council to OMEA (311723-GENEPAD) and in part by an advanced investigator grant from the European Research Council to SHR (787270 - Paget-Advance). The PRISM trial was supported by grants from the Arthritis Research Campaign (13627) and the Paget’s Association.

